# All three MutL complexes are required for repeat expansion in a human stem cell model of CAG-repeat expansion mediated glutaminase deficiency

**DOI:** 10.1101/2023.12.26.573357

**Authors:** Bruce Hayward, Daman Kumari, Saikat Santra, Clara D.M. van Karnebeek, André B.P. van Kuilenburg, Karen Usdin

## Abstract

The Repeat Expansion Diseases (REDs) arise from the expansion of a disease-specific short tandem repeat (STR). Different REDs differ with respect to the repeat involved, the cells that are most expansion prone and the extent of expansion. Furthermore, whether these diseases share a common expansion mechanism is unclear. To date, expansion has only been studied in a limited number of REDs. Here we report the first studies of the expansion mechanism in induced pluripotent stem cells derived from a patient with a form of the glutaminase deficiency disorder known as Global Developmental Delay, Progressive Ataxia, And Elevated Glutamine (GDPAG; OMIM# 618412) caused by the expansion of a CAG-STR in the 5’ UTR of the glutaminase (*GLS*) gene. We show that alleles with as few as ∼120 repeats show detectable expansions in culture despite relatively low levels of R-loops formed at this locus. Additionally, using a CRISPR-Cas9 knockout approach we show that PMS2 and MLH3, the constituents of MutLα and MutLγ, the 2 mammalian MutL complexes known to be involved in mismatch repair (MMR), are essential for expansion. Furthermore, PMS1, a component of a less well understood MutL complex, MutLβ, is also important, if not essential, for repeat expansion in these cells. Our results provide insights into the factors important for expansion and lend weight to the idea that, despite some differences, the same mechanism is responsible for expansion in many, if not all, REDs.

## Introduction

Global Developmental Delay, Progressive Ataxia, And Elevated Glutamine (GDPAG; OMIM# 618412; aka Glutaminase Deficiency, GLSD) is a severe neurological disorder resulting from mutations in the *Glutaminase* (*GLS*) gene, a gene encoding the enzyme required for conversion of glutamine to glutamate, a critical excitatory neurotransmitter in the brain. Recently it has been demonstrated that some GLSD cases are caused by expansion of a CAG-tandem repeat tract located in the 5’ untranslated region of the *Glutaminase* (*GLS*) gene ^1^. More than 50 other human genetic disorders have been identified that also result from expansions of short tandem repeats (STRs) or microsatellites ^2^. These disorders are known collectively as the Repeat Expansion Diseases (REDs).

The REDs are caused by different STRs. They also differ in the cell types most prone to expansion and the extent of repeat expansion, with the number of repeats added varying widely from just a few repeats in the case of Huntington’s disease (HD) to hundreds or even thousands of repeats in the case of other disorders such as Myotonic Dystrophy Type 1 (DM1) ^2^. Whether the same mechanism is responsible for expansions in all REDs is still the subject of debate. While no study of the expansion mechanism in GLSD has been reported to date, expansion has been studied in models of other REDs including HD, DM1, the Fragile X related disorders (FXDs) and Friedreich’s ataxia (FRDA), disorders involving expansion of CAG, CTG, CGG and GAA-STRs respectively. Unlike the microsatellite instability (MSI) associated with cancer predisposition, mouse and cell models of the REDs suggest that some mismatch repair (MMR) proteins are actually required for expansion in these diseases ^3–5^, and recent genome wide association studies (GWAS) and analysis of single nucleotide polymorphisms (SNPs) have shown that some of these same factors are modifiers of both somatic expansion risk and disease severity in patient cohorts ^6–9^.

The MMR factors known to be important for expansion in most model systems include the MutSβ mismatch recognition complex and MutLγ, one of the two MLH1-binding complexes known to be involved in mismatch processing in MMR ^10^. We had previously shown that in addition to MutLγ, a heterodimer of MLH1 and MLH3, that the second mismatch processing complex, MutLα, a heterodimer of MLH1 and PMS2, was also required for expansion in a mouse embryonic stem cell (mESC) model of the FXDs ^11^. Furthermore, MutLβ, a heterodimer of MLH1 and PMS1, a relatively abundant complex whose function is unclear, is also important, perhaps essential for expansion in these cells ^11^. The fact that all three MutL complexes are important for expansion was surprising since it suggested a hitherto unknown cooperation between these factors. Furthermore, an essential role for PMS2 was inconsistent with the demonstration that in a mouse model of DM1, loss of PMS2 resulted in a loss of only ∼50% of expansions in the somatic tissues examined ^12^, while in a mouse model of FRDA, PMS2 actually suppressed expansions ^13^. This raised the question of whether the results we obtained in the mouse ESC model system were disease relevant not only for the FXDs, but for other REDs as well.

To begin to address this question, we made GLSD patient-derived iPSCs and showed that the large disease-associated alleles, as well as a contracted allele that arose spontaneously during iPSC derivation, all expand in culture. This contracted allele provided us with a useful model system to study the expansion process in this disease. We then used a gene knockout approach to examine the role of PMS1, PMS2 and MLH3 in these cells. This allowed us to establish not only whether these factors promote or protect against expansion but whether they are essential for either process. This is important since an essential contribution indicates that these complexes cooperate to generate the expansions rather than acting redundantly. We show here that all 3 MLH1-binding partners are also all important for expansions in the GLSD patient-derived iPSCs, with PMS2 and MLH3 both being essential and with PMS1 also playing a vital, perhaps essential, role. This has implications not only for our understanding of the expansion process but for the possibility of a common therapeutic approach to reducing repeat expansion in multiple REDs as well.

## Materials and Methods

### Reagents

All reagents were from Sigma-Aldrich (St. Louis, MO) unless otherwise specified. Primers were from Life Technologies (Grand Island, NY). Capillary electrophoresis of fluorescently labeled PCR products and Sanger sequencing of plasmid constructs was carried out by Psomagen (Rockville, MD). All restriction enzymes were from New England Biolabs (Ipswich, MA). Plasmid pSpCas9(BB)-2A-Puro (PX459) V2.0 was a gift from Feng Zhang (Addgene plasmid 62988; http://n2t.net/addgene:62988) and carries Cas9 from *Streptococcus pyogenes* ^14^. Plasmid pCK002_U6-Sa-sgRNA(mod)_EFS-SaCas9-2A-Puro_WPRE was a gift from Aviv Regev (Addgene plasmid 85452; http://n2t.net/addgene:85452) and carries Cas9 from *Staphylococcus aureus* ^15^. pCE-mp53DD, carrying a dominant-negative p53 fragment that improves cell survival after CRISPRing, was a gift from Shinya Yamanaka (Addgene plasmid 41856; http://n2t.net/addgene:41856) ^16^.

### Plasmid construction

pXSaCas9-Purov2 was created by excising the gRNA scaffold, the SpCas9 promoter and the SpCas9 ORF of pX459v2 with *Bbs*I + *Bst*EII digestion and replacing them with an equivalent PCR fragment containing the SaCas9 gRNA scaffold, the SaCas9 promoter and the SaCas9 ORF from Addgene plasmid 85452 using a Gibson Assembly strategy ^17^. Sanger sequencing was performed to ensure the fidelity of the inserted PCR product. p53DD-alone was derived from pCE-mp53DD by *Hind*III + *Bam*HI digestion and recircularization to remove the EBNA1 and oriP sequences to prevent potential episomal propagation of the plasmid. Suitable gRNAs with minimal predicted off-targets for *MLH3* and *PMS1* were identified using the Synthego knockout guide algorithm (https://design.synthego.com/<u>#/</u>). Because of the complex nature of the *PMS2* locus ^18^, no acceptable SpCas9 gRNAs could be identified using either the Synthego algorithm or the Benchling (benchling.com) prediction tools. However, using the CRISPOR suite (http://crispor.tefor.net/, ^19^) a suitable gRNA for SaCas9 that had minimal predicted off-targets was identified. Oligonucleotide pairs encoding these gRNAs were synthesized using MLH3_gRNA-F (CACCGTTGCTGTTGATGTAAGCAG) and MLH3_gRNA-R (AAACCTGCTTACATCAACAGCAAC); PMS1_gRNA-F (CACCGACATTTTATAGGAGAACTA) and PMS1_gRNA-R (AAACTAGTTCTCCTATAAAATGTC) and PMS2_gRNA-F (CACCGTCAAACCGTACTCTTCACACA) and PMS2_gRNA-R (GAACTGTGTGAAGAGTACGGTTTGAC) for *MLH3*, *PMS1* and *PMS2* respectively.

The oligonucleotide pairs were annealed and cloned into either pX459v2 (*MLH3* and *PMS1*) or pXSaCas9-Purov2 (*PMS2*) which had been digested with *Bbs*I. All gRNA-encoding plasmids were verified by Sanger sequencing.

### Cell culture

The derivation of iPSCs was carried out by the NHLBI iPSCs core facility using the Cytotune 2.0 Sendai virus kit (Thermo Fisher Scientific, Waltham, MA, A16517) with a few modifications for high throughput reprogramming essentially as described earlier ^20^. Fibroblasts from an individual carrying a CAG expansion in the 5’-UTR of the GLS locus on both alleles (proband P2 in ^1^) were used as the cell source. Pluripotency was assessed by staining for Tra-160, SSEA4 and Nanog by FACS analysis as described earlier ^20^. The iPSC cells were routinely maintained feeder-free in StemFlex media (Thermo Fisher Scientific, A3349401) in wells coated with laminin-521 (StemCell Technologies, 200-0117). To passage the cells StemPro Accutase (Thermo Fisher Scientific, A11105-01) was used to dissociate them and, after transfer to a fresh well the StemFlex was supplemented with RevitaCell (Thermo Fisher Scientific, A2644501) for 24 hours. The iPSCs were cultured at 37°C until they were Sendai Virus (SeV)-free as assessed by RT-qPCR with a FAM-labeled SeV Taqman assay (Thermo Fisher Scientific, assay ID Mr04269880_mr; 4331182) and VIC-labeled *hGUSB* Taqman probe/primers (Thermo Fisher Scientific, 4326320E) as the normalizing control. Using the same methods, Fragile X premutation (FXPM) iPSCs with 147 CGG repeats were derived from a male FX fibroblast cell line mosaic for different sized *FMR1* alleles.

### DNA:RNA immunoprecipitation (DRIP) assay

DNA was isolated from cells using the salting out method ^21^. DRIP was performed in duplicate for DNAs from FXPM iPSCs and in triplicate for DNAs from GLSD iPSCs using the anti RNA:DNA hybrid antibody (S9.6, Millipore-Sigma, MABE1095). For each DNA sample, three DRIPs were performed: normal mouse IgG (Millipore-Sigma, 12-371), S9.6 antibody without RNAse H treatment and RNAse H treatment followed by S9.6 antibody. A total of 25 µg DNA was either mock digested or digested with 30 units of RNAse H (New England Biolabs, M0297) in 100 µl final volume at 37 °C for 6 hours. The volume was made up to 400 µl with 300 µl of ChIP dilution buffer (0.01% SDS, 1.1% Triton X-100, 1.2 mM EDTA, 16.7 mM Tris-HCl, pH 8.1, 167 mM NaCl) and sonicated to fragments <500 bp using Bioruptor® (6 minutes at medium setting, 30 seconds ON/30 seconds OFF). To 350 µl of the sonicated DNA, 650 µl of the ChIP dilution buffer and 10 µl of protease inhibitor cocktail (Sigma-Aldrich, P8340) was added and mixed. An aliquot (1%) was saved as input sample. The sonicated DNA was then precleared with 50 µl of Protein A agarose beads/Salmon sperm DNA slurry (EMD Millipore, 16–157) for one hour on a rotator in cold. The precleared supernatant was incubated with 5 µg of mouse IgG or S9.6 antibody overnight on a rotator in a cold room. The sample was then incubated with 60 µl of the Protein A agarose beads/Salmon sperm DNA slurry to collect the immune complexes. The material was washed once for 5 minutes each with low salt wash buffer (0.1% SDS, 1% Triton X-100, 2 mM EDTA, 20 mM Tris-HCl, pH 8.1, 150 mM NaCl), high salt wash buffer (0.1% SDS, 1% Triton X-100, 2 mM EDTA, 20 mM Tris-HCl, pH 8.1, 500 mM NaCl), LiCl wash buffer (0.25 M LiCl, 1% IGEPAL-CA630, 1% deoxycholic acid (sodium salt), 1 mM EDTA, 10 mM Tris, pH 8.0), and twice with TE pH 8.0 (10 mM Tris-HCl, 1 mM EDTA, pH 8.0). The immunoprecipitated material was eluted from the beads using 2 x 250 µl of elution buffer (1% SDS, 100 mM NaHCO3). The input and DRIP samples were then treated with phenol/chloroform and precipitated overnight at −20 °C with 0.3 M sodium acetate and ethanol in the presence of pellet paint. After washing with 70% ethanol, the samples were resuspended in 50 µl 0.1X TE pH 8.0. Real-time PCRs on the immunoprecipitated DNAs were carried out in triplicate in 20 µl final volume using the PowerUP^TM^ SYBR™ Green PCR master mix (Thermo Fisher Scientific, A25742) and 2 µl of DNA using QuantStudio^TM^ 3 real-time PCR system (Thermo Fisher Scientific). For the amplification of *GLS* promoter region upstream of the repeat, 150 nM each of primer GLS-F2 (5’ GATTTGAGCCAATCGCAGC 3’) and GLS-R1 (5’ GGCTAGAGACCTTCAGCGCT 3’) were used and for the amplification of *GLS* exon1 region downstream of the repeat, 300 nM each of primer GLS Ex1-F (5’-cccaagtagctgccctttcc-3’) and GLS Ex1-R (5’-cgctcaacaggggaggatg-3’) were used. For the amplification of positive control *FMR1* exon1 region, 300 nM each of *FMR1* Exon1-F (5’-CGCTAGCAGGGCTGAAGAGAA-3’) and *FMR1* Exon1-R (5’-GTACCTTGTAGAAAGCGCCATTGGAG-3’) were used. For the amplification of negative control region *ZNF554*, 300 nM each of *ZNF554*-F (5’-CGGGGAAAAGCCCTATAAAT-3’) and *ZNF554*-R (5’-TCCACATTCACTGCATTCGT-3’) were used.

### RNA quantitation

Total RNA was isolated from cells using TRIzol^TM^ reagent (Thermo Fisher Scientific, 15596026) and quantified on DS-11 Spectrophotometer (DeNovix, Wilmington, DE). Three hundred nanograms of total RNA was reverse transcribed in 20 µl final volume using SuperScript^TM^ IV VILO^TM^ master mix (Thermo Fisher Scientific, 11766050) as per manufacturer’s instructions. Real-time PCR was performed in triplicate using 2 µl of the cDNA, FAM-labeled *FMR1* (Hs00924547_m1), FAM-labeled *ZNF554* (Hs01014440_m1), FAM-labeled *GLS* (Hs01014020_m1) and VIC-labeled *β-glucoronidase* (*GUSB*) endogenous control (4326320E) Taqman probe-primers (Thermo Fisher Scientific) and TaqMan® FAST universal PCR master mix (Thermo Fisher Scientific, 4444556) using a StepOnePlus^TM^ Real-Time PCR system (Thermo Fisher Scientific). For quantitation, the comparative threshold (Ct) method was used.

### Generation of CRISPR knockouts

To generate the desired CRISPR knockouts ∼100K iPSCs were plated into a 12-well 24 hours before transfection. Immediately prior to transfection the media was replaced with 0.5 ml of OptiMem (Thermo Fisher Scientific, 31985062) supplemented with RevitaCell. Transfections were carried out using 800 ng of the appropriate Cas9 plasmid and 200 ng of p53DD-alone complexed with 4μl of Lipofectamine Stem (Thermo Fisher Scientific, STEM00008) as per the manufacturer’s instructions. Four hours after the transfection, 0.5 ml of StemFlex supplemented with RevitaCell was added to the cells. Approximately 24 hours after the transfection, the media was replaced with 1 ml StemFlex supplemented with RevitaCell containing 1 μg/ml Puromycin and selection applied for 24-48 hours. The culture was allowed to recover for 4 days and then ∼4K cells were plated into a 60 mm dish to allow single cell derived colonies to appear. Twenty-four colonies were picked, genomic DNA prepared using the KAPA Express reagent (a component of the KAPA Mouse Genotyping Kit from Roche, 07961804001) and analyzed for both CRISPR lesions and repeat size as described below.

The lesions present in the *MLH3* and *PMS1* lines were identified by direct sequencing of the PCR product containing the gRNA target and subsequent analysis by the ICE tool from Synthego (ice.synthego.com). Sequencing of the cDNA was used to confirm the loss of the entire target exon in one *MLH3* clone that did not generate a genomic PCR product (MLH3 #3). The lesions present in the individual *PMS2* cell lines were identified by sequencing of individual plasmid clones derived from the PCR product containing the gRNA target. A known SNP (rs1805319) allowed the 2 alleles to be unambiguously identified. The mutations generated are shown in Table 1. The absence of the targeted protein was confirmed for PMS1 and PMS2 by western blotting using standard protocols (Fig. S1). However, two different MLH3 antibodies tested showed no specific bands corresponding to MLH3 even in a control cell line (HepG2) and since the cell lines showed no DNA evidence of a functional *MLH3* allele, no additional antibodies were tested. Loss of any one of the MLH1-binding partners had no significant effect on the levels of either PMS1 or PMS2 (Fig. S1). Since no good MLH3 antibody was available, the effects on MLH3 could not be assessed. However, evidence in mice and humans suggests that a decrease in the level of MLH3 would also not be expected from the loss of either PMS1 or PMS2 ^11,22,23^.

### Analysis of repeat size

The GLS repeats in the GLSD B cell line were amplified using the primer pair GLS-F and GLS-R, as previously described ^1^ except that GLS-R was 5’-FAM labelled. The PCR reaction (15 µl) contained 50 mM Tris-HCl pH 9.0, 1.75 mM MgCl_2_, 22 mM (NH_4_)_2_SO_4_, 0.5 µM each primer, 1.5 M betaine, 2% DMSO and 0.2 mM each dATP, dCTP, dGTP and dTTP. Three tenths of a microliter of Robust HotStart KAPA2G polymerase (Roche, 07961316001) was added per 100 μl of PCR mix. Cycling parameters of 98°C for 3 minutes, 35 cycles of (98°C for 30 seconds, 65°C for 30 seconds, 72°C for 210 seconds), and 72°C for 10 minutes were used. The *FMR1* repeats in the FXPM iPSC line were amplified as previously described ^24^. Both *GLS* and *FMR1* PCR products were then sized by agarose gel and capillary electrophoresis. The fragment analysis data files from the capillary electrophoresis were analyzed using a custom R script as previously described ^25^. The FXPM line has 147 repeats at *FMR1* and is homozygous for a normal *GLS* allele, while the GLSD B line had one allele of ∼120 repeats and another of ∼900 repeats at *GLS* and carried a normal *FMR1* allele (data not shown).

### Statistical methods

Statistical significance of observed differences was calculated using paired two-tailed Student’s t test (GraphPad Software, Inc., La Jolla, CA), and a p-value of ≤0.05 was considered statistically significant. Specific details can be found in the results and the figure legends.

## Results

### Repeat expansions are seen in GLSD patient derived iPSCs

We derived iPSCs from fibroblasts from an individual with expansions in both *GLS* alleles (Patient P2 in ^1^). These fibroblasts contained a heterogeneous mixture of alleles of different sizes and, as a result, iPSC lines with a range of different repeat sizes were obtained, having ∼120 to ∼900 repeats in each allele. As can be seen from PCR analysis of the repeat over time in culture, ongoing expansions like those reported in iPSCs from other REDs ^6,26–28^ could be detected by agarose gel electrophoresis (Fig. S2). Specifically, the alleles in the population appear to expand in concert, a phenomenon consistent with the high frequency addition of a small number of repeats ^29^. While large alleles are not well resolved by agarose gel electrophoresis, thus precluding an accurate determination of repeat number, we estimate that larger alleles in the GLSD C and GLSD D lines gained approximately 60 repeats over a period of 50-59 days (Fig. S2), corresponding to the gain of ∼1 repeat/day or ∼1 expansion event/day in the majority of cells in the population.

Most of these alleles are too large to be analyzed by capillary electrophoresis (CE), a simple cost-effective, high-resolution technique that allows alleles differing by one repeat to be reliably distinguished ^25^. However, one iPSC line (GLSD B) had sustained a contraction resulting in an allele with ∼120 repeats, putting it in the range suitable for CE analysis. We then derived additional cell lines from this one by plating the iPSC line for single colonies. This resulted in individual cell lines with repeat numbers that varied by 1-4 repeats. Six of these lines were grown for 49 days. As can be seen from Fig. 1 and Fig. 2, the allele in two of the 6 lines gained 7 repeats over 49 days in culture as evidenced by a clear shift of the modal allele by 7 repeats.

**Fig. 1.**
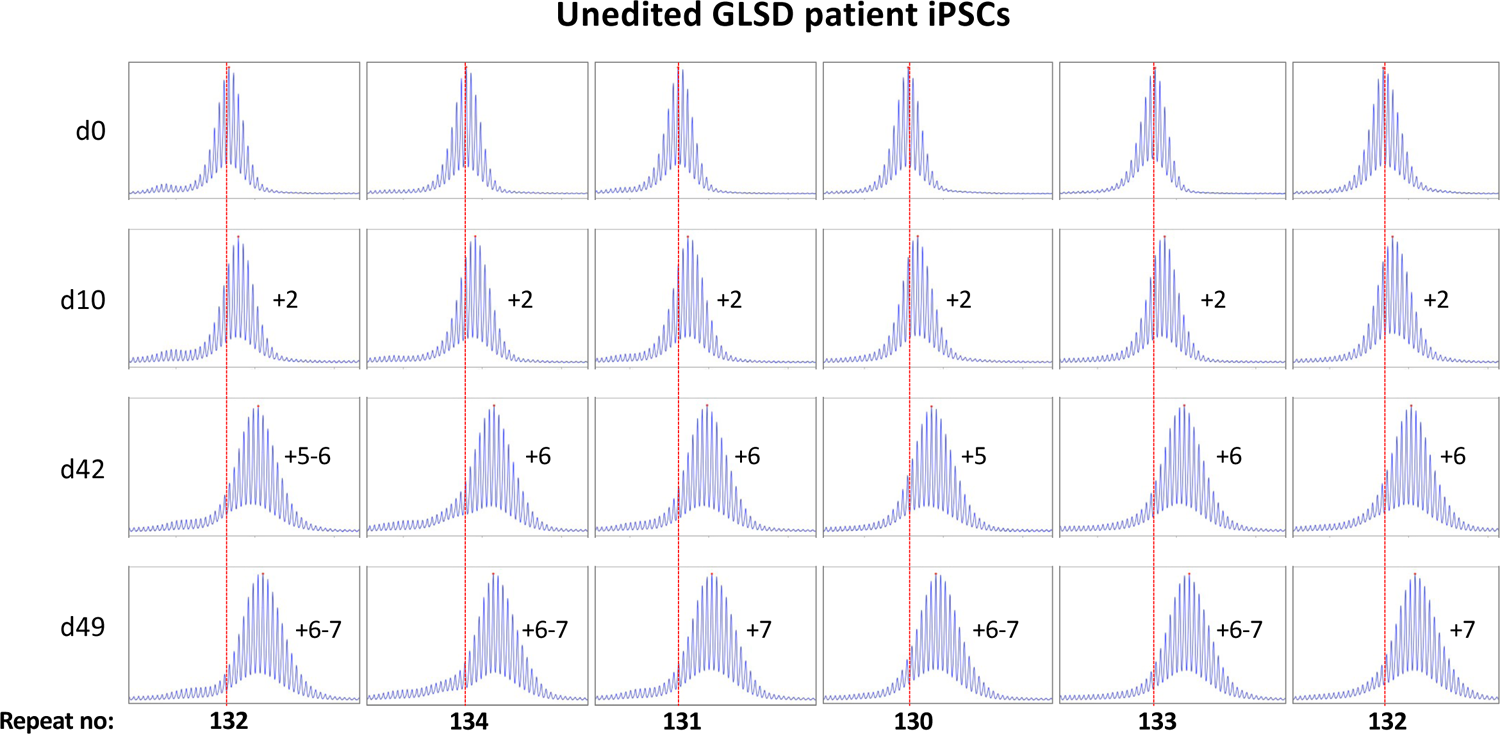
Repeat PCR profiles of 6 clones derived from unedited patient iPSCs. Six clones with initial modal repeat numbers of 130-134 were grown for 49 days and samples removed for analysis at the indicated time points. In each case, the red dotted line indicates the repeat number present in the original culture. The numbers alongside each plot indicate the change in repeat number at that time-point.

**Fig. 2.**
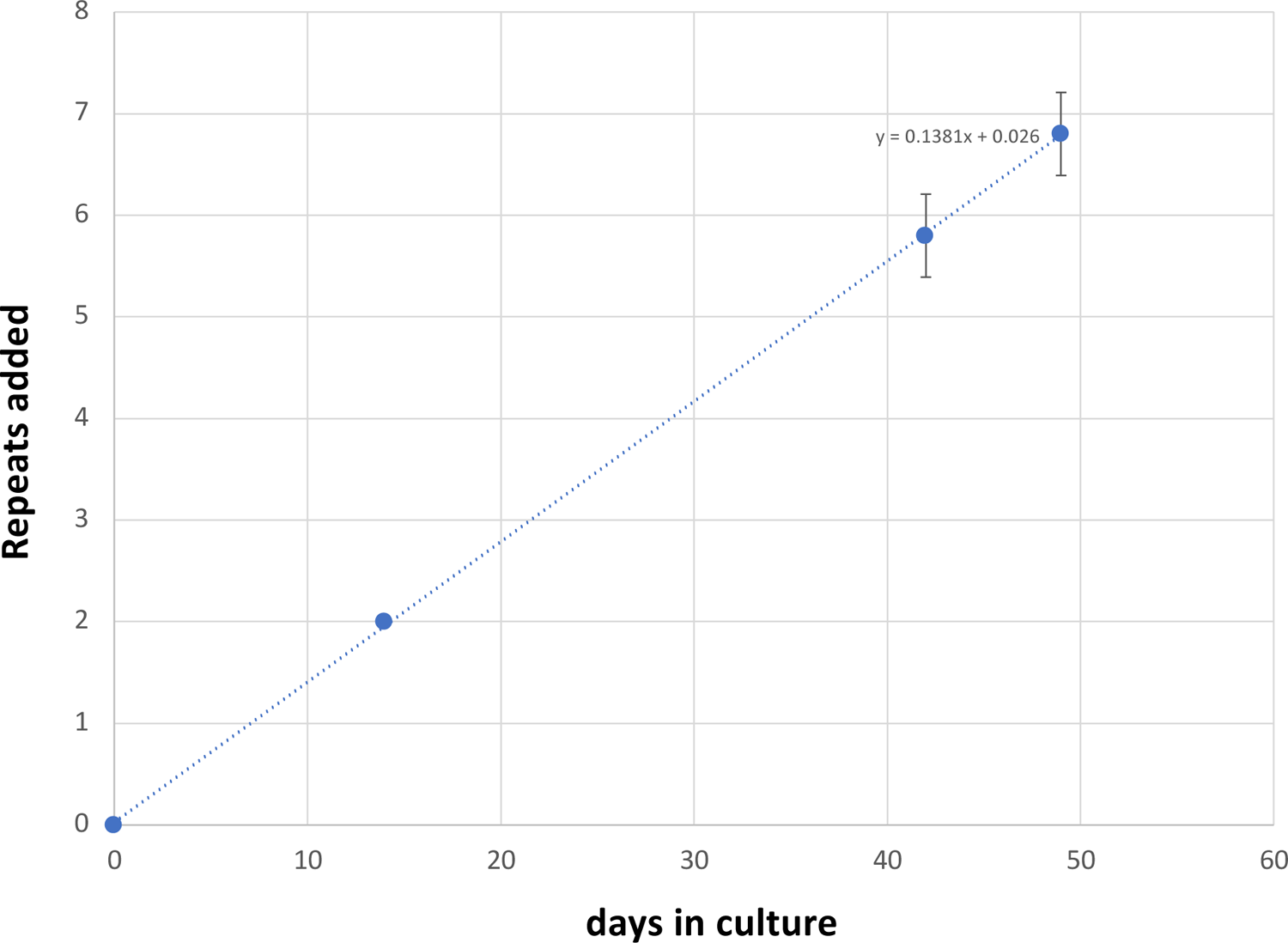
Graphical representation of the average change in repeat number over time in the unedited patient iPSC lines shown in. Fig. 1. The change in the modal repeat number for each cell line at each time point was averaged and plotted along with the standard deviation for each point. The equation for the regression line through the data points is shown.

In the case of the remaining cell lines the two largest allele populations were very similar in abundance making it impossible to identify a single modal allele. Since these allele sizes corresponded to a gain of 6 and 7 repeats respectively, it suggests that these cell lines gained on average ∼6.5 repeats over the 49 days in culture. Thus, these lines are all expanding at a rate of ∼0.14 repeats/day or ∼ 1 repeat/week.

### R-loop formation at the GLS locus

R-loops, three-stranded nucleic acid structures containing an RNA:DNA hybrid, are seen at some disease-associated STRs, where they have been suggested to play a role in the expansion process ^10,30–33^. To explore this idea in the context of *GLS* expansion, we used the S9.6 antibody ^34^ to assess the levels of R-loop formed in FXPM iPSCs lacking a *GLS* expansion and the GLSD iPSCs using *ZNF554* and *FMR1* as negative and positive controls respectively ^35–38^. As can be seen in Fig. 3a, both of the iPSC lines produced a similar DRIP signal at the *GLS* promoter and exon 1 of the gene, even though GLS mRNA levels in the GLSD iPSCs are more than 4-fold lower (Fig. 3b). The DRIP signals seen at both *GLS* regions tested were significantly lower than the DRIP signal at *FMR1* in both the FXPM and GLSD iPSCs and not significantly different from the DRIP signals seen for the negative control in these lines. Since the R-loop-negative control gene, *ZNF554,* and *GLS* are expressed at comparable levels in GLSD iPSCs (Fig. 3b), the expanded GLS locus does not meet the criteria for a stable R-loop forming region.

**Fig. 3.**
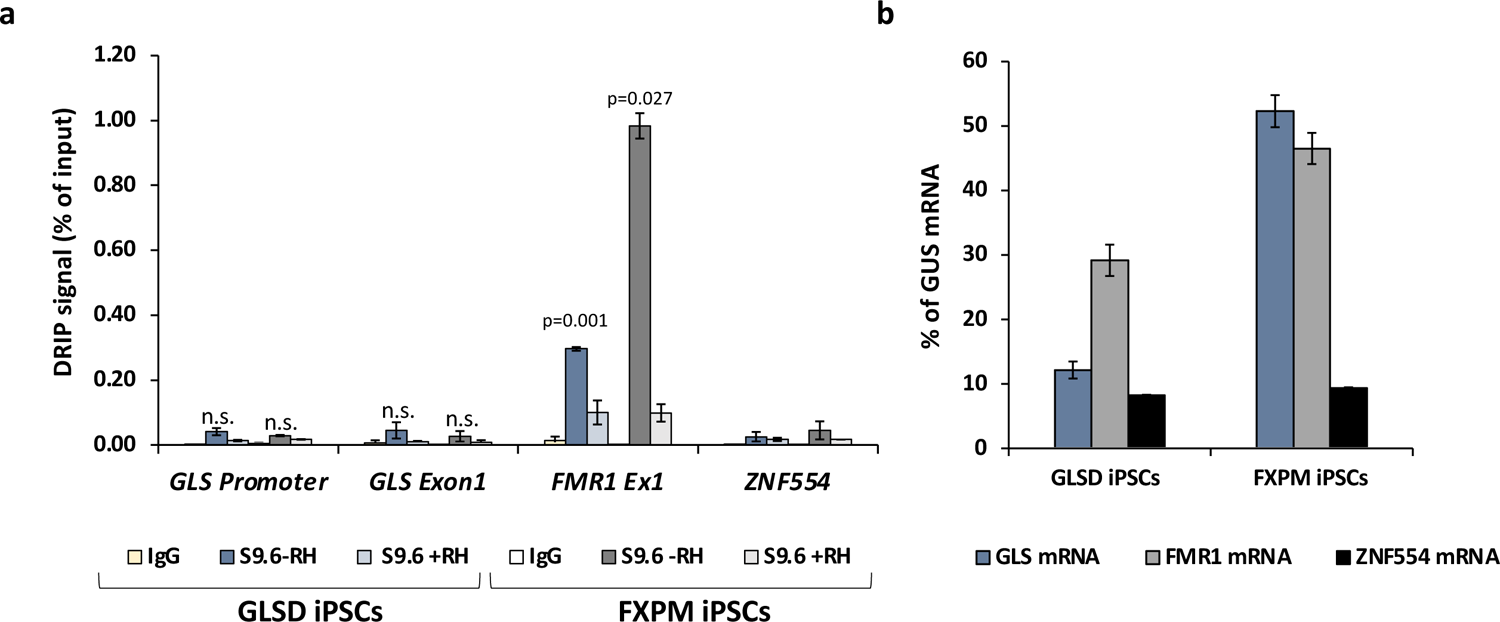
DNA:RNA immunoprecipitation (DRIP) signal at the *GLS* promoter and exon 1 region in unaffected control and patient iPSCs. Data shown is an average of two and three independent experiments for FXPM iPSCs and GLSD iPSCs respectively. Error bars represent standard deviation. The *FMR1* exon 1 was used as a positive control and the *ZNF554* gene was used as a negative control. Two-tailed, paired student t test was used to calculate p-values. The p-values shown indicate the comparison with the negative control (*ZNF554*). n.s., not significant.

### Null mutations in PMS1, PMS2 and MLH3 eliminate almost all expansions in the GLSD iPSCs

Using CRISPR-Cas9 we made null mutations in *PMS1*, *PMS2* and *MLH3* as described in the Materials and Methods. We isolated three independent sequence-verified null lines for each gene. The mutations generated in each cell line are listed in Table 1. In the case of the *PMS1* null lines, two lines had repeat numbers within the range of the unedited lines, with one cell line having one repeat fewer than the smallest unedited line. All of the *PMS2* and *MLH3* null lines had repeat numbers within the range of the unedited lines. We then grew each of these lines in culture for 49 days (Fig 4 – 6).

**Fig. 4.**
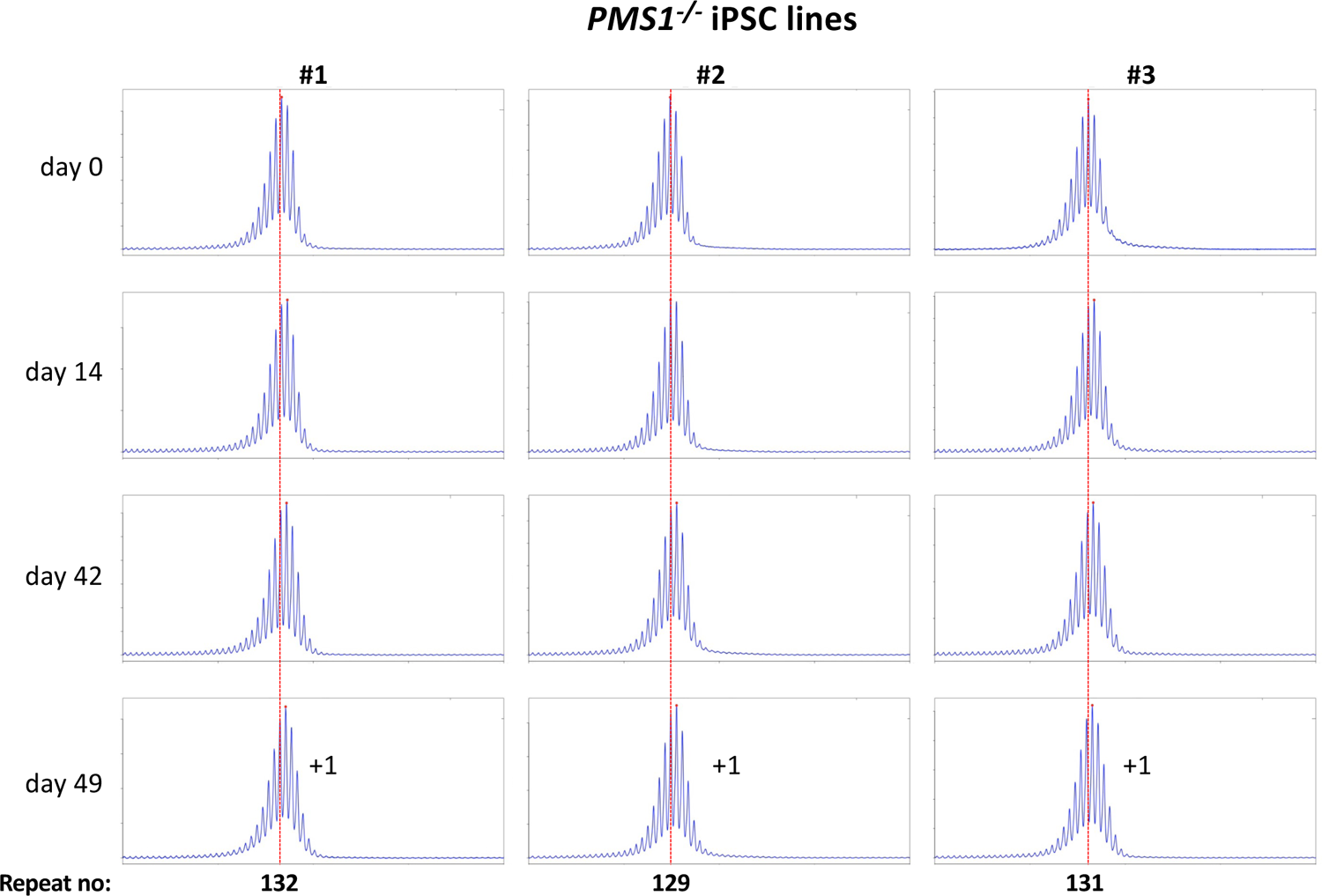
Repeat PCR profiles of 3 different *PMS1^-/-^* patient iPSC lines. Three lines with similar repeat numbers were grown for 49 days and samples removed for analysis at the indicated time points. In each case, the red dotted line indicates the repeat number present in the original culture. The numbers alongside the last plot indicate the change in repeat number at that time-point.

As can be seen from Fig. 4, each of the *PMS1* null lines showed a subtle shift in their PCR profile that resulted in the modal allele being one repeat larger than the modal allele in the original culture. This represents at least an 86% reduction in the extent of expansion compared to unedited lines. The number of repeats in the *PMS2* and *MLH3* null lines did not change over the same period (Fig 5 - 6), consistent with the idea that MutLα and MutLγ are both required for expansion in these cells.

**Fig. 5.**
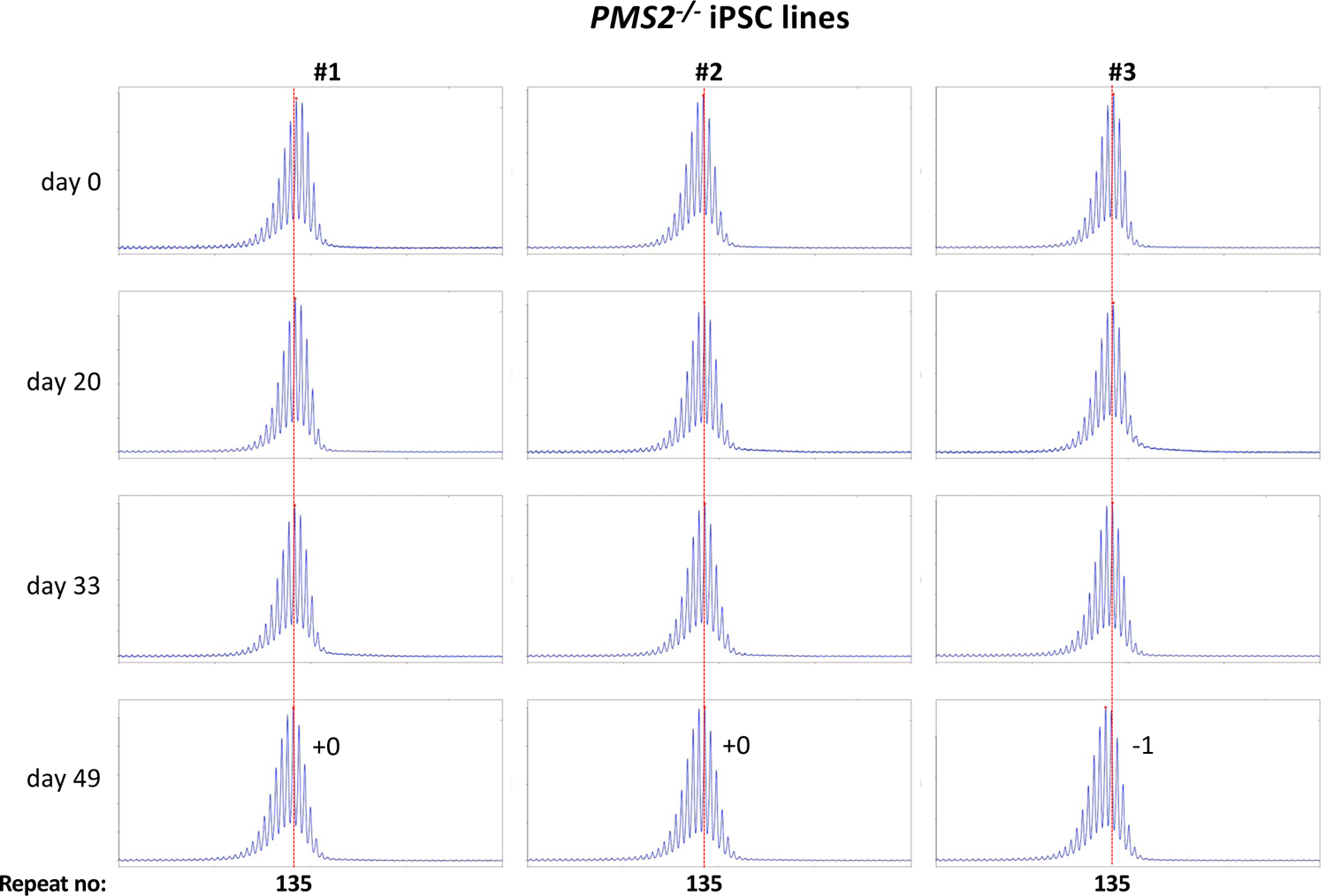
Repeat PCR profiles of 3 different *PMS2^-/-^* patient iPSC lines. Three lines with similar repeat numbers were grown for 49 days and samples removed for analysis at the indicated time points. In each case, the red dotted line indicates the repeat number present in the original culture. The numbers alongside the last plot indicate the change in repeat number at that time-point.

**Fig. 6.**
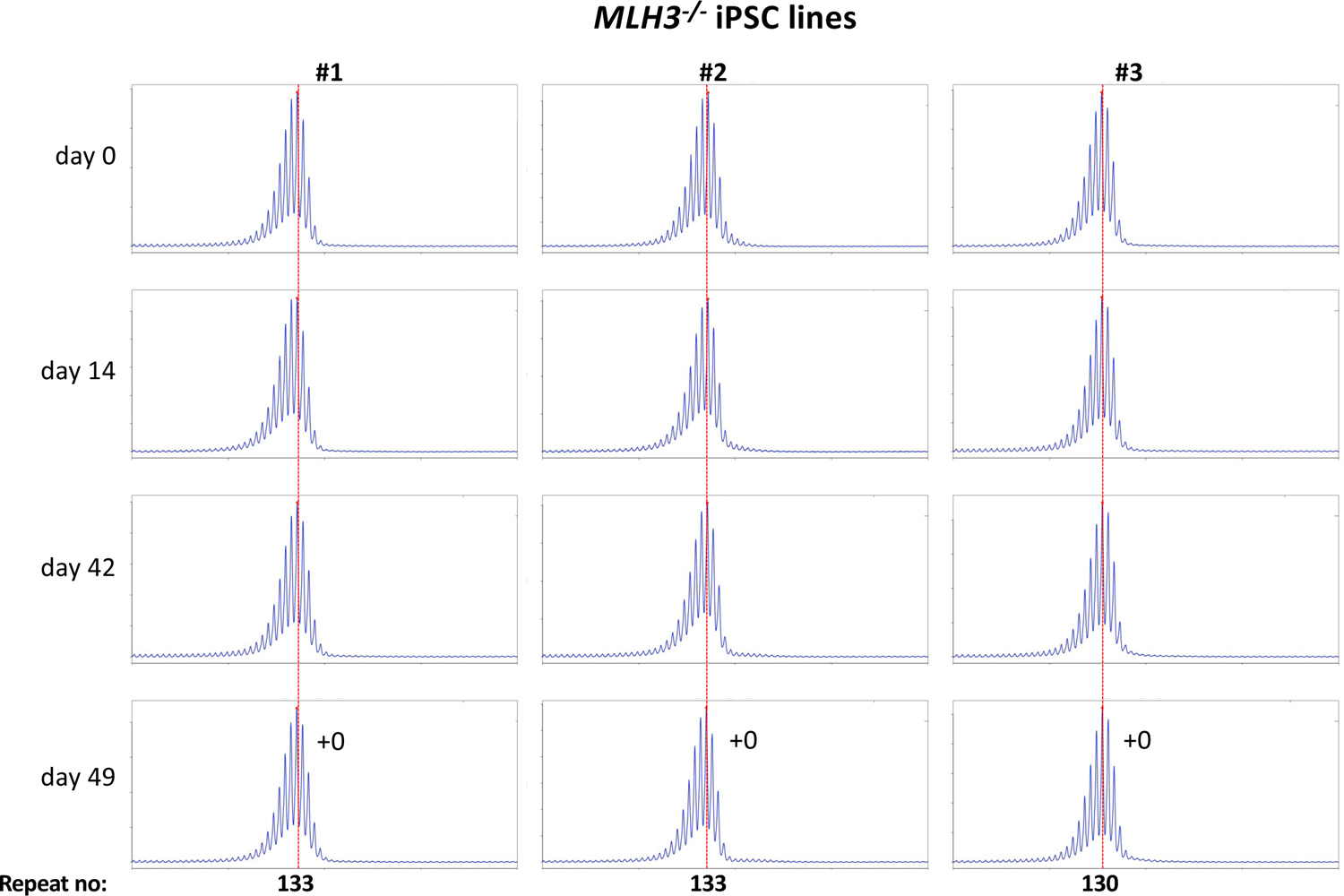
Repeat PCR profiles of 3 different *MLH3^-/-^* patient iPSC lines. Three lines with similar repeat numbers were grown for 49 days and samples removed for analysis at the indicated time points. In each case, the red dotted line indicates the repeat number present in the original culture. The numbers alongside the last plot indicate the change in repeat number at that time-point.

## Discussion

Here we demonstrate that GLSD patient-derived iPSCs constitute a tractable human model system to study “expansion in a dish”, with alleles as small as ∼120 repeats showing clear evidence of expansion over an experimentally reasonable timeframe. This smaller allele can be resolved by capillary electrophoresis thus allowing the accurate measurement of changes in repeat number at single repeat resolution and the ready evaluation of factors that impact expansion.

However, while we saw expansion in these cells (Figs 1-2), we saw little evidence of significant amounts of stable R-loop formation in cells heterozygous for the small allele and an allele with ∼900 repeats (Fig. 3). This suggests that such R-loops may not play a major role in promoting expansion in GLSD. This is consistent with the observation that many expansion-prone repeats show a strong AT-skew rather than the GC-skew thought to be important for R-loop formation ^39,40^. However, transcription is required for expansion in many REDs ^4^, and our data does not preclude a contribution from the RNA:DNA hybrid formed as part of normal transcription. Hybrid formation may create a window of opportunity for out-of-register realignment of the repeats behind the transcription complex. This could result in the formation of the first loop-out in the expansion substrate illustrated in the model shown in Fig. S3. However, additional experiments are required to definitively address the contribution of R-loops to the expansion process.

We also used these cells to examine the effect of CRISPR-generated null mutations in *PMS1*, *PMS2* or *MLH3*, genes we had previously shown to be required for expansions in mESCs from a mouse model of the FXDs ^11^. The effect of the loss of each of these proteins was broadly similar to what we found in the mESC model ^11^. Specifically, in the case of *PMS2* and *MLH3*, no repeats were gained. This is consistent with the idea that both of these MLH1-binding partners are essential for expansion in these cells. However, in contrast to the FXD mESCs where 1-2 repeats were lost over a similar period, no repeats were lost in this case. We had previously attributed the loss of repeats in the FXD mESCs to the effect of the MLH3 or PMS2-deficiency on normal MMR. The failure to see a similar loss in the GLSD-patient iPSCs may reflect the smaller number of repeats in these lines. This might make the repeat tract less prone to any microsatellite instability resulting from an MMR-deficiency. As we saw with the FXD mESCs, loss of PMS1 resulted in the growth of the repeat tract being limited to a single repeat, in this case corresponding to an 86% decrease in expansions over a 7-week period. Whether the residual repeat gained reflects a small fraction of PMS1-independent expansion events, or some effect of the absence of PMS1 on normal MMR is unclear.

Our results clearly demonstrate that the contribution of all three MutL proteins to expansion is not unique to the mouse model of the FXDs. Rather it may be a general feature of expansion in the REDs. This raises the question of whether PMS2’s paradoxical effects, namely its ability to protect against expansions in some model systems and its requirement for expansion in others, may not reflect different mechanisms for generating expansions in different diseases but rather differences related to cell type and/or the sequence and/or location of the repeat in the genome. For example, as illustrated in the model shown in Fig. S3, it is possible that PMS2’s promutagenic and antimutagenic effects on expansion are related to the different cleavage preferences of MutLα and MutLγ ^41,42^ combined with the relative levels of MLH3 and PMS2 in the cell type being evaluated.

Our finding that PMS1 is also involved in generating expansions in GLSD lends weight to the idea that PMS1 is an important determinant of expansion risk in multiple REDs. Since loss of PMS1 is not associated with increased tumor risk in mice ^43^ and its role in human MMR is unclear ^44^, it may thus be a useful target for therapeutic development.

## Data availability statement

All data generated or analyzed during this study are included in this published article and its Supplementary Information files.

## Acknowledgements

We thank the patient and family for providing the cells used in this study. We would also like to acknowledge Carson Miller for his help with weekend media changes. We thank Flora Tassone for the human fibroblast line from which iPSCs with FXPM alleles were derived.

## Supplementary Methods

### Western blotting

To prepare total cell lysates, cells were pelleted at 250 x g for 5 min and washed once with ice-cold PBS supplemented with 1X protease inhibitor cocktail (Sigma-Aldrich, St. Louis, MO, P8340). The cell pellet was then resuspended in the lysis buffer (10 mM Tris Cl pH 7.5, 1 mM EDTA pH 8.0, 1% Triton X-100, 1X protease inhibitor cocktail) and incubated on ice for 10 min followed by sonication (3 cycles of 30 seconds ON/30 seconds OFF) at medium setting using Bioruptor® (Diagenode, Denville, NJ) to solubilize proteins and shear the DNA. The protein amount was quantified using Bio-Rad Protein Assay Dye Reagent Concentrate (Bio-Rad Laboratories, Inc., Hercules, CA, #5000006) as per manufacturer’s protocol. Before using the lysate for western blot analyses, 1X volumes of Novex® LDS Sample Buffer (Thermo Fisher Scientific, Waltham, MA, NP0007) and NuPAGE® Sample Reducing Agent (Thermo Fisher Scientific, NP0009) were added, and the samples were heated at 75 °C for 10 min.

Twenty (for PMS1) or thirty (for PMS2) micrograms of total cell lysates were run on NuPAGE^TM^ 4-12% Bis-Tris gel (Thermo Fisher Scientific, NP0322BOX) and transferred to nitrocellulose membrane using the Trans-Blot Turbo RTA mini 0.2 µm nitrocellulose transfer kit (1704270) and Trans-Blot® Turbo^TM^ Transfer system from Bio-Rad Laboratories. The membrane was blocked for two hours with 5% blocking agent (GE Healthcare Bio-Sciences, Pittsburg, PA, RPN2125) in TBST (1X Tris buffered saline with 0.1% Tween 20). The following antibodies were used, 1:1000 diluted anti-PMS1 rabbit monoclonal antibody [EPR27158-78] (Abcam, ab315798) overnight at 4°C, 1:500 diluted anti-PMS2(B3) mouse monoclonal antibody (Santa Cruz Biotechnology, # sc-25315) overnight at 4°C, 1:10,000 diluted anti-b-actin mouse monoclonal antibody (Invitrogen # MA1-140) for 1 hour at room temperature. Following incubation with the primary antibodies, the blots were washed three times with TBST for 5 minutes each and then probed with either 1:2000 diluted HRP-labeled secondary Rabbit antibody (Millipore Sigma, (GENA934) or 1:5000 diluted HRP-labeled secondary mouse antibody (Millipore Sigma (12-349) for 1 hour. The blot was then washed three times with TBST for 5 minutes each and once with TBS. The signal was detected using ECL^TM^ Prime detection reagents (GE Healthcare Bio-Sciences) and imaged with ChemiDoc imaging system (Bio-Rad Laboratories).

### Supplementary Figures

**Fig. S1.**
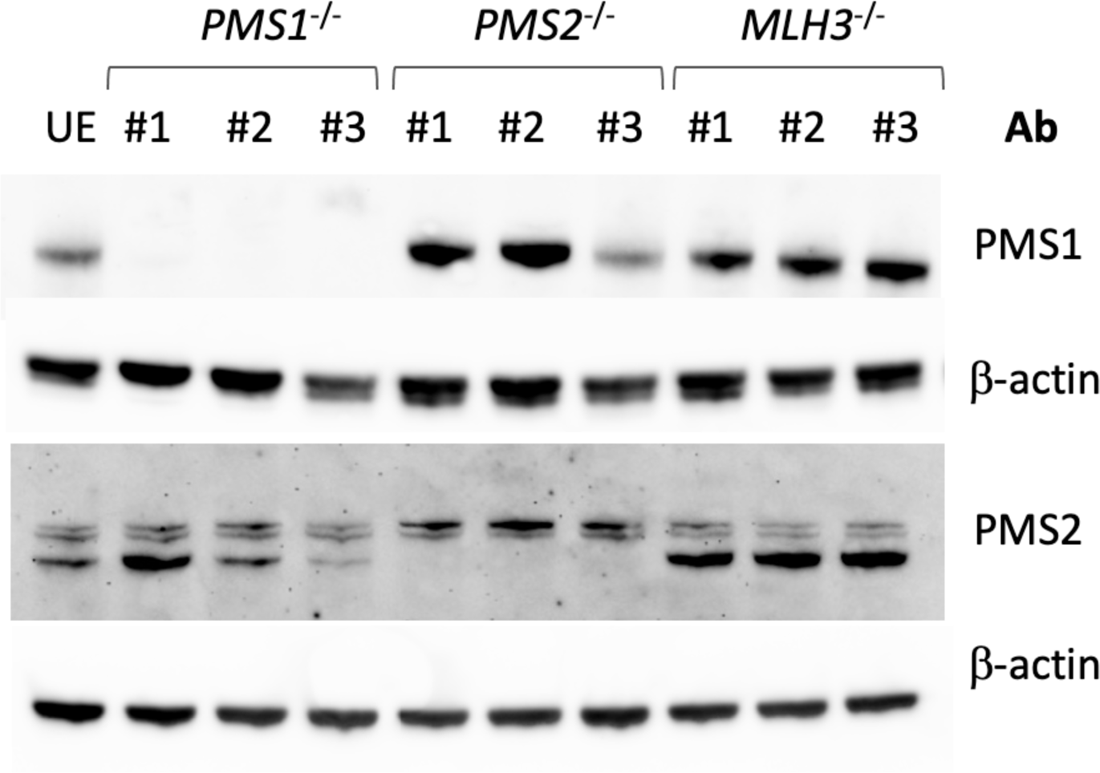
Analysis of MLH1-binding proteins in PMS1, PMS2 and MLH3 gene edited lines. a) Western blots of the unedited (UE) GLSD patient iPSC line and the *PMS1*, *PMS2* and *MLH3* null lines derived from it used in this study challenged with antibodies for PMS1, PMS2 and β-actin.

**Fig. S2.**
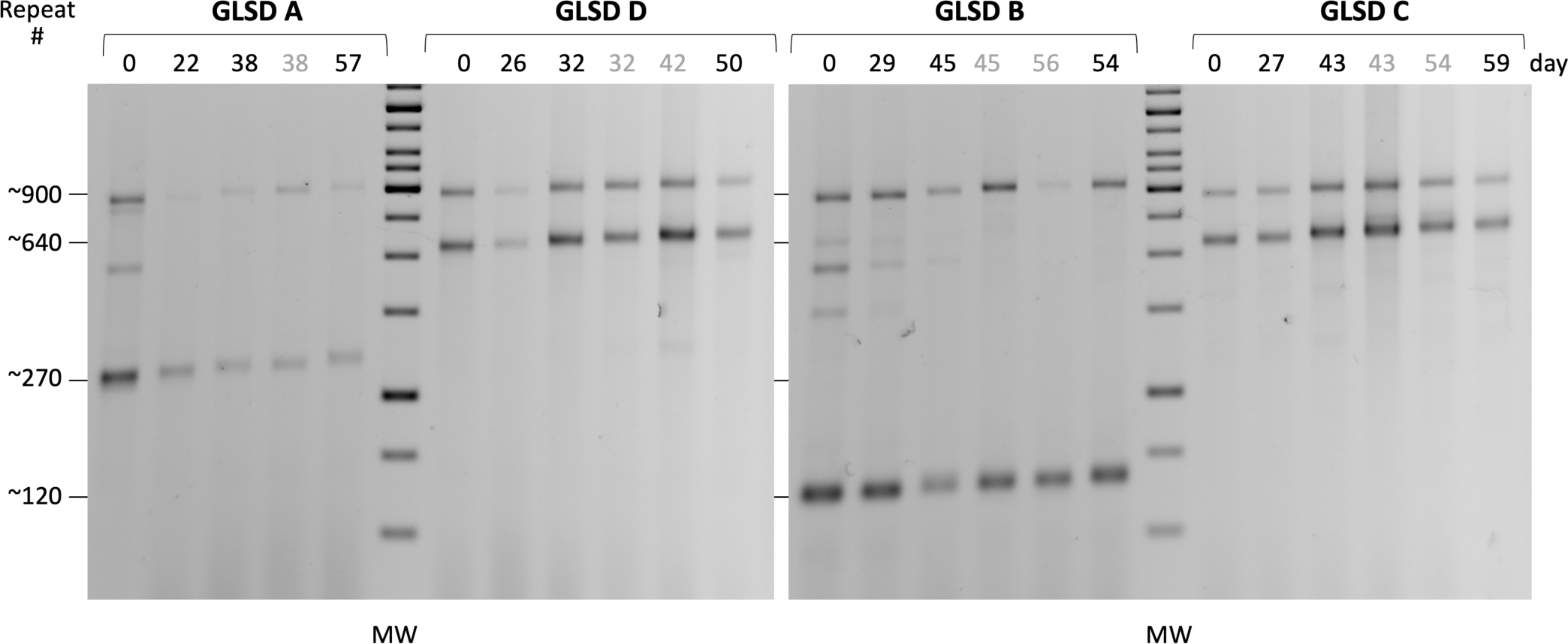
Expansion of the CAG-repeat tract over time in 4 different patient iPSC lines. The iPSC lines were propagated in culture as described in the Materials and Methods section and DNA isolated from the lines at different time points as indicated. The PCR products obtained by amplification through the GLS repeat in iPSCs derived from a patient (Patient 2 ^1^) were then resolved by agarose gel electrophoresis alongside a GeneRuler 1 kb DNA ladder (Thermo Fisher Scientific, SM0311). The approximate repeat numbers in different starting alleles are indicated alongside. The samples shown in grey font were obtained from cultures that had been grown at 38.5 °C for 6 days prior to harvesting to eliminate residual Sendai virus. Interestingly, they show a slightly more expansion than the cells maintained at 37°C which may point to an effect of temperature on expansion rate.

**Fig. S3.**
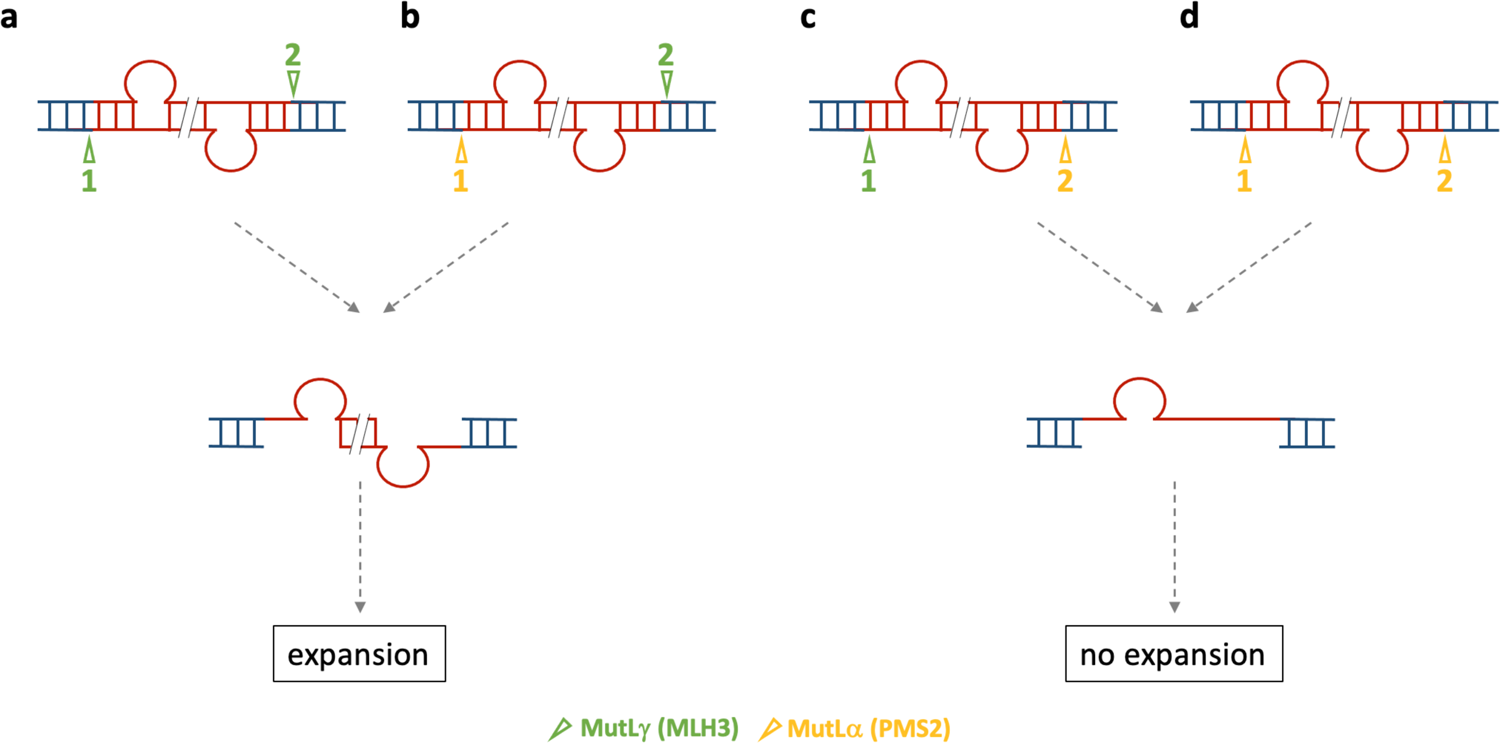
Model for PMS2’s promutagenic and antimutagenic roles in repeat expansion. In this model, the substrate for expansion is a double loop-out structure formed in the repeat. Formation of the substrate during transcription may explain the transcriptional dependence ^2^. It may also explain the high frequency with which expansion occurs. The size of the loop-outs determines the net gain of repeats and thus the size of the expansion. Since some expansion-prone repeats do not form hairpins and most expansions involve the addition of a single repeat, a loop-out of a single repeat is shown. However, it may be that larger loop-outs form, with or without hairpin formation, and this could contribute to the small number of larger expansions that are occasionally seen. Since PMS1 has no nucleolytic activity ^3^, it is presumed to have a structural role in expansion that is not shown in this model. Each loop-out is bound by MutLγ or MutLα and the binding depending on some combination of the relative abundance of the two complexes and their relative binding affinities. Each MutL protein then makes a cut on one or other strand, depending on their cleavage preferences, with MutLγ cutting the strand opposite the loop-out and MutLα showing no bias for which strand is cut when there is no pre-existing nick and cutting on the nicked strand when there is ^4,5^. Each possible set of cuts is shown assuming that the first cut (cut 1) is performed by the MutL complex bound to the bottom strand loop-out. When both cuts (cuts 1 and 2) are made by MutLγ (a), an intermediate containing gaps opposite each loop-out is produced. Secondary structures formed by the looped-out bases might limit further resection that would otherwise result in the loss of one or both of the looped-out regions. Filling of both gaps by a gap-filling polymerase and ligation results in an expansion. Gap-filling of each loop-out adds a total of 3 bases on each strand, resulting in a net gain of one repeat. If MutLα makes the first cut and it occurs on the strand opposite the loop-out to which it is bound (b) and there is enough MutLγ to make the second cut, then it too will result in an expansion. On the other hand, if MutLγ makes the first cut and MutLα the second (c), or if MutLα makes both cuts (d) this would result in two cuts on the same strand. Repair synthesis in any of these cases would then restore the original allele. Thus, in cells where MutLγ is sufficient to process all the expansion substrates (a), they would all be processed to generate expansions. When MutLα predominates, all substrates would be processed so as to restore the original allele (d), and when MutLγ and MutLα are both involved in processing the expansion substrate, some would be processed to generate expansions (b) and some fraction would be processed so as to restore the original allele (c) Since the expansion rate depends on the rate at which the substrate forms and is processed, rates that would be shared amongst all cells of a given type, all cells in a homogenous population of cells would expand at a similar rate. This results in a similar shift of most of the population to progressively larger alleles with time.

## Original Images

**Fig. S1.**
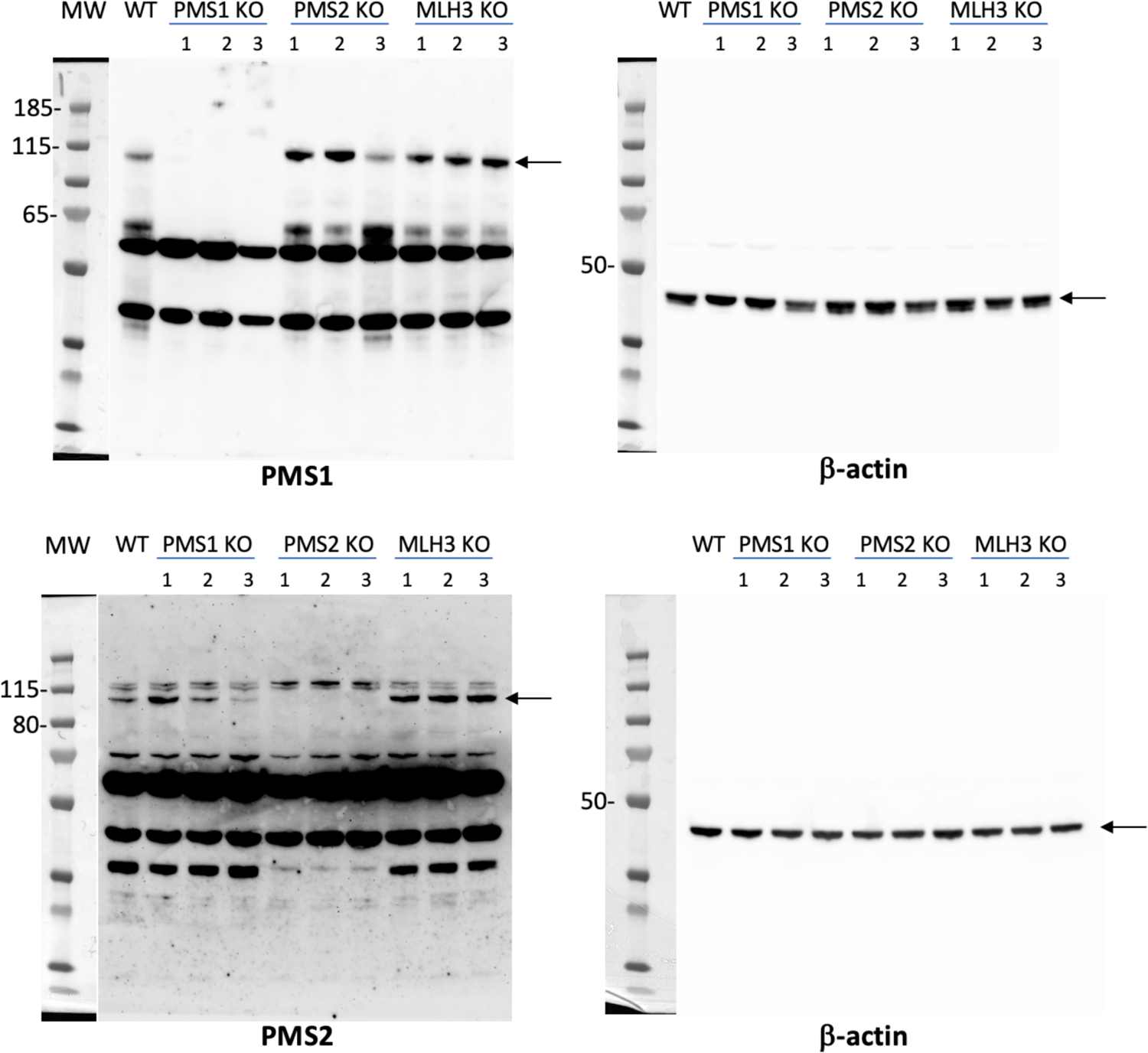

**Fig. S2.**
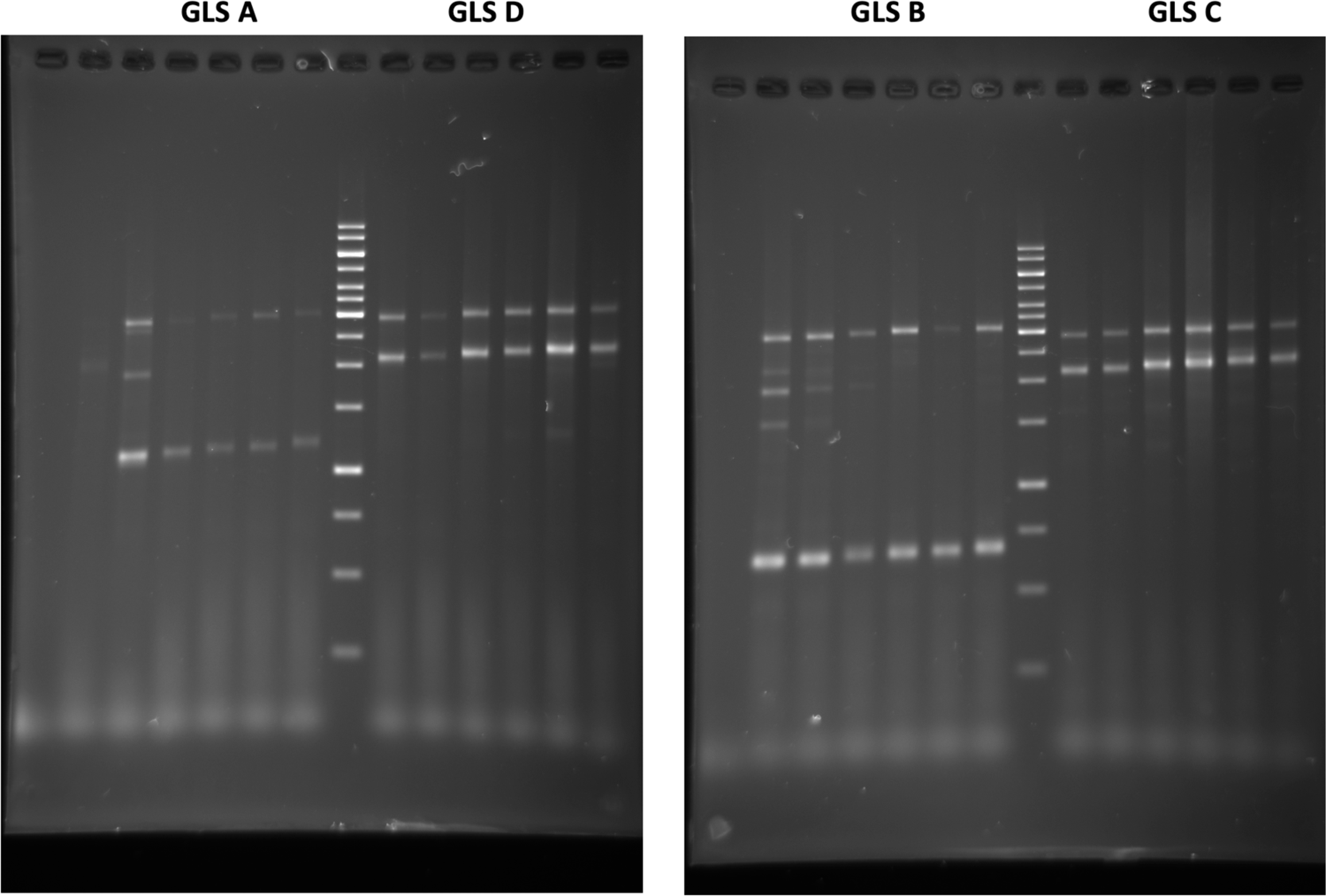

